# Niacin fine-tunes energy homeostasis through canonical GPR109A signaling

**DOI:** 10.1101/382416

## Abstract

Niacin has long been considered as a high-potency drug for beneficially treating lipid abnormalities, however, its anti-atherosclerotic effects have been challenged by recent studies. Here, we demonstrated that oral supplementation of niacin resulted in a significant reduction in body weight and fat mass without affecting food intake in high-fat diet-fed wild-type mice, but not in GPR109A-defeicient mice. Further investigation showed that niacin challenge led to a remarkable inhibition of hepatic lipogenesis via a GPR109A-dependent ERK1/2/AMPK pathway. Additionally, we demonstrated that niacin treatment stimulated thermogenesis in brown adipose tissue by induction of thermogenic genes via GPR109A. Moreover, we observed that mice exposed to niacin exhibited a dramatic decrease in intestinal absorption of fatty acids. Together, our data demonstrate that acting on GPR109A, niacin shows the potential to maintain energy homeostasis by fine-tuning hepatic lipogenesis, BAT/beige thermogenesis and intestinal fat absorption, representing a potential approach to the treatment of lipid abnormalities.

## Introduction

The incidence of overweight and obesity has become a global epidemic, with over 1.9 billion overweight and 600 million obese individuals worldwide (Ng et al, 2014; Wang et al, 2011). Obesity is caused by an abnormal excess storage of energy as lipids in adipose tissue due to a net imbalance between energy intake and energy expenditure (Lowell & Spiegelman, 2000; Olsen et al, 2017). Obesity is considered a major risk factor for numerous comorbidities including major diseases such as cardiovascular disease, diabetes, and cancer (Haslam & James, 2005; Kopelman, 2000). However, despite the recent tremendous efforts, effective antiobesity pharmacotherapies are still limited. Additionally, to date, current antiobesity agents are designed to reduce food-intake and appetite through the hypothalamus-based molecular pathways, but unfortunately, these agents exhibit severe psychiatric or cardiovascular side effects (Cunningham & Wiviott, 2014; Manning et al, 2014; Moreira et al, 2009). Therefore, the magnitude of this current health epidemic has heightened the need for alternative therapeutic strategies aiming to control body weight through potential targets in peripheral non-neuronal tissues.

Since discovery of decreasing plasma cholesterol levels in 1955, nicotinic acid (niacin) has been used to treat dyslipidemia as an approved drug for more than half a century (Altschul et al, 1955; Carlson, 2005). A number of randomized clinical trials demonstrated that niacin monotherapy has been shown to substantially increase levels of HDL-C and decrease levels of triglycerides (Elam et al, 2000; Guyton et al, 1998). However, its exact mechanism was not well understood until the orphan receptor GPR109A (recently renamed as hydroxyl-carboxylic acid receptor 2 or HCA2) was identified as a receptor with high affinity for niacin in 2003, independently by three research groups (Offermanns et al, 2011; Soga et al, 2003; Tunaru et al, 2003; Wise et al, 2003). Subsequently, the ketone body β-hydroxy butyrate was identified as an endogenous ligand for GPR109A (Taggart et al, 2005). In adipocytes, upon stimulation by niacin, GPR109A functions via Gi/o proteins to lower intracellular cAMP production, leading to decreased protein kinase A-mediated activation of hormone-sensitive lipase (HSL), which in turn reduces triglyceride hydrolysis to free fatty acids (FFA) (Digby et al, 2009). This FFA hypothesis is supported by evidence that niacin failed to lower FFA and TGs in GPR109A-knockout mice (Tunaru et al, 2003).

The established clinical benefits of niacin treatment on cardiovascular events have been challenged by recent studies showing that niacin exerts its beneficial effects on GPR109A-mediated anti-atherosclerotic activity independently from its anti-lipolytic properties (Lukasova et al, 2011a; Wu et al, 2010). However, these data did not exclude the possibility that niacin provides cardiovascular benefits in an anti-lipolysis-independent manner through GPR109A. Niacin-mediated beneficial cardiovascular effect has been shown to be transferable with GPR109A-competent bone marrow cells (Wu et al, 2010). In addition, accumulating evidence has demonstrated that activation of GPR109A was able to induce anti-inflammatory effects not only on atherosclerosis, also on acute ischemic stroke, arthritis, chronic renal failure and sepsis through inhibition of immune cell chemotaxis and pro-inflammatory cytokine production (Chen et al, 2014;

Cho et al, 2009; Digby et al, 2012; Kwon et al, 2011; Lukasova et al, 2011a; Mitrofanov et al, 2005; Rahman et al, 2014; Shehadah et al, 2010; Shi et al, 2017; Wu et al, 2010). Therefore, more studies are needed to thoroughly evaluate the GPR109A-mediated pleiotropic effects on atherogenesis and other metabolic diseases.

GPR109A-induced inhibition in adipocyte lipolysis and fatty acid mobilization in response to long-term treatment with niacin would result in excessive synthesis of TGs and obesity (Ganji et al, 2004). However, during breeding process of GPR109A-knockout mice we introduced from Dr. Offermanns’ lab, we found that the GPR109A-knockout mice are more prone to being obese compared to the wild-type mice. Therefore, we initiated this study to investigate roles of niacin/GPR109A in the regulation of lipid metabolism using GPR109A-knockout mice combined with high fat diet (HFD)-induced obesity mouse model. We show that upon activation by niacin, GPR109A fine-tunes lipid metabolism through multipathways including inhibition of hepatocyte lipogenesis and fatty acid absorption and promotion of brown adipose tissue (BAT) thermogenesis.

## Results

### GPR109A deficient animals display obesity

To explore GPR109A function in lipid metabolism, parallel experiments in mice with GPR109A (wild-type) and without GPR109A (Gpr109a^−/−^) were performed. When both wild-type and Gpr109a^−/−^ male mice were fed with either high-fat diet (HFD) or normal-chow diet (NCD) for 12 weeks, we recorded weight gain weekly. As the feeding progressed, starting from approximately 10-12 weeks of age, the GPR109A-knockout mice exhibited significantly increased body weight compared with the wild-type mice (Figs 1A and C, and EV1A and C). In addition, we monitored food intake weekly throughout experimental period, if any, not significant difference in food intake between GPR109A-knockout and wild-type mice was observed (Figs 1B and EV 1B). After 12 weeks of normal-chow diet feeding, adipose tissue weights and organ weights were examined. Our data demonstrated that GPR109A-deficient mice displayed a significant increase in adipose tissue weights including epididymal, perirenal and inguinal white adipose tissue and liver weight, but not in BAT or other organ weights (kidney, lung, heart and spleen) when compared to wild-type mice (Figs 1E and EV1D). Moreover, H&E stained histological sections of eWAT and electron microscope observation of liver and muscle revealed that GPR109A-deficient mice had a striking visible increase in amount and size of lipid droplets in liver, muscle and eWAT compared with the wild-type group (Fig 1D).

**Figure 1.**
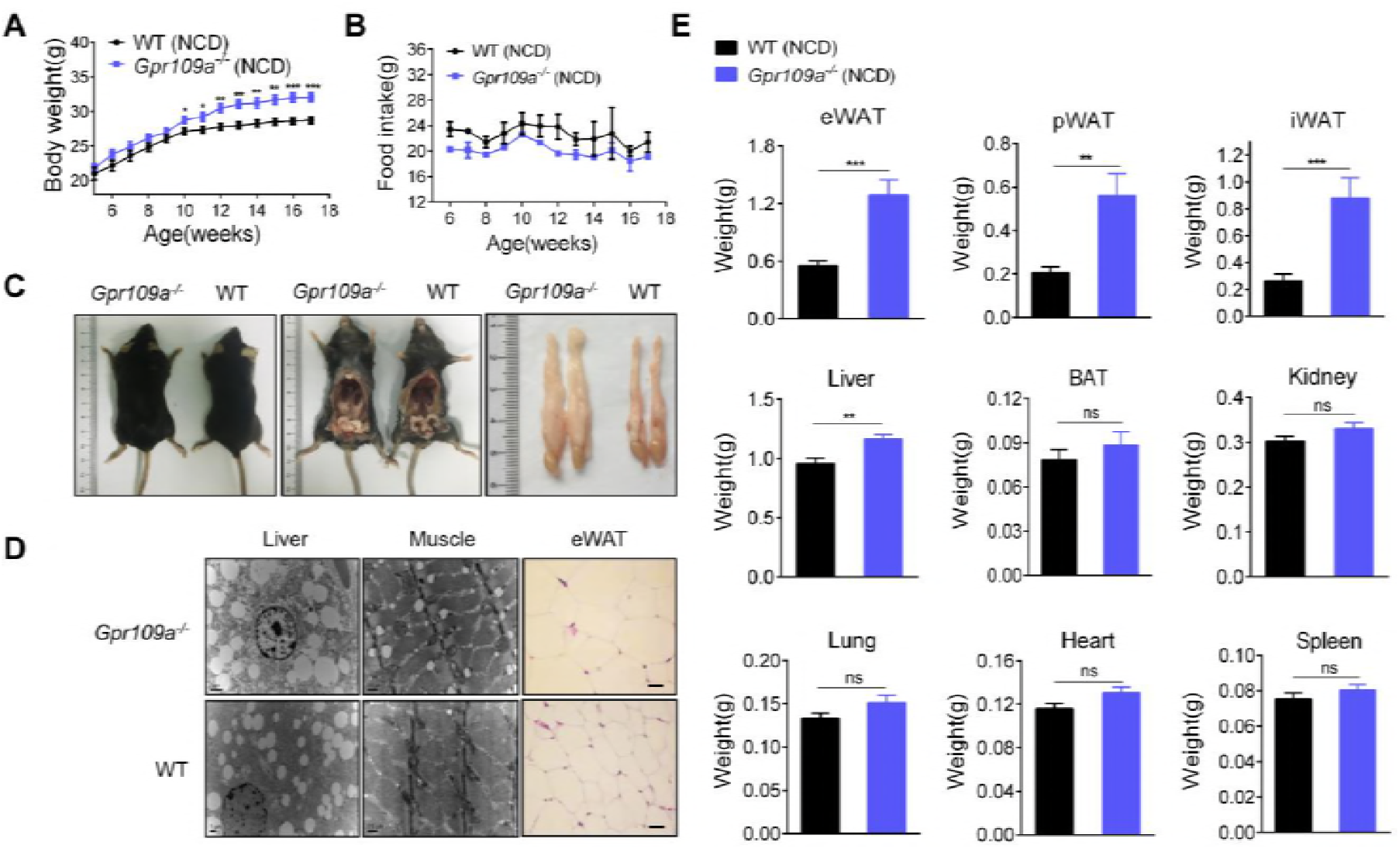
GPR109A deficient animals display obesity. A,B Body weight (A) and food intake (B) were recorded weekly (n=10-11). C Representative images of fat mass were shown. D Representative SEM images in liver (left panels; scale bars, as shown in figure) and muscle (middle panels; scale bars, as shown in figure), and H&E staining of epididymal white adipose tissue (eWAT) (right panels; scale bars, 100 μm). E Fat pads and organ weights were recorded after mice anesthetized (n=10-11). Data information: All the data are presented as means ±SEM. In (A, B, E), data were analyzed by unpaired two-tailed student’s t-test. \**P* < 0.05; \*\**P* < 0.01; \*\*\**P* < 0.001; \*\*\*\**p* < 0.0001.

### Niacin exerts inhibitory effects on HFD-induced obesity

To assess the effect of niacin on lipid metabolism, wild-type mice were fed with a 6-week high-fat diet (HFD) to induce obesity. HFD supplemented with niacin (HFD + niacin, 500 mM) or vehicle (HFD + water) was then treated during the following 8 weeks. As anticipated, the mice maintained on the HFD exhibited a progressive increase in body weight gain. Treatment with niacin resulted in a significant reduction in body weight, but weekly food intake remained unchanged as compared to vehicle-treated mice (Fig 2A, B and C). Mice fed with HFD in response to challenge with niacin exhibited a significant reduction in adipose tissue weights including epididymal, perirenal and inguinal white adipose tissue and liver weight, but not in BAT or other organ weights (kidney, lung, heart and spleen) with respect to vehicle-treated mice (Fig 2E). Histological examination showed that niacin challenge led to a marked decrease in lipid accumulation in liver, muscle and eWAT (Fig 2D).

**Figure 2.**
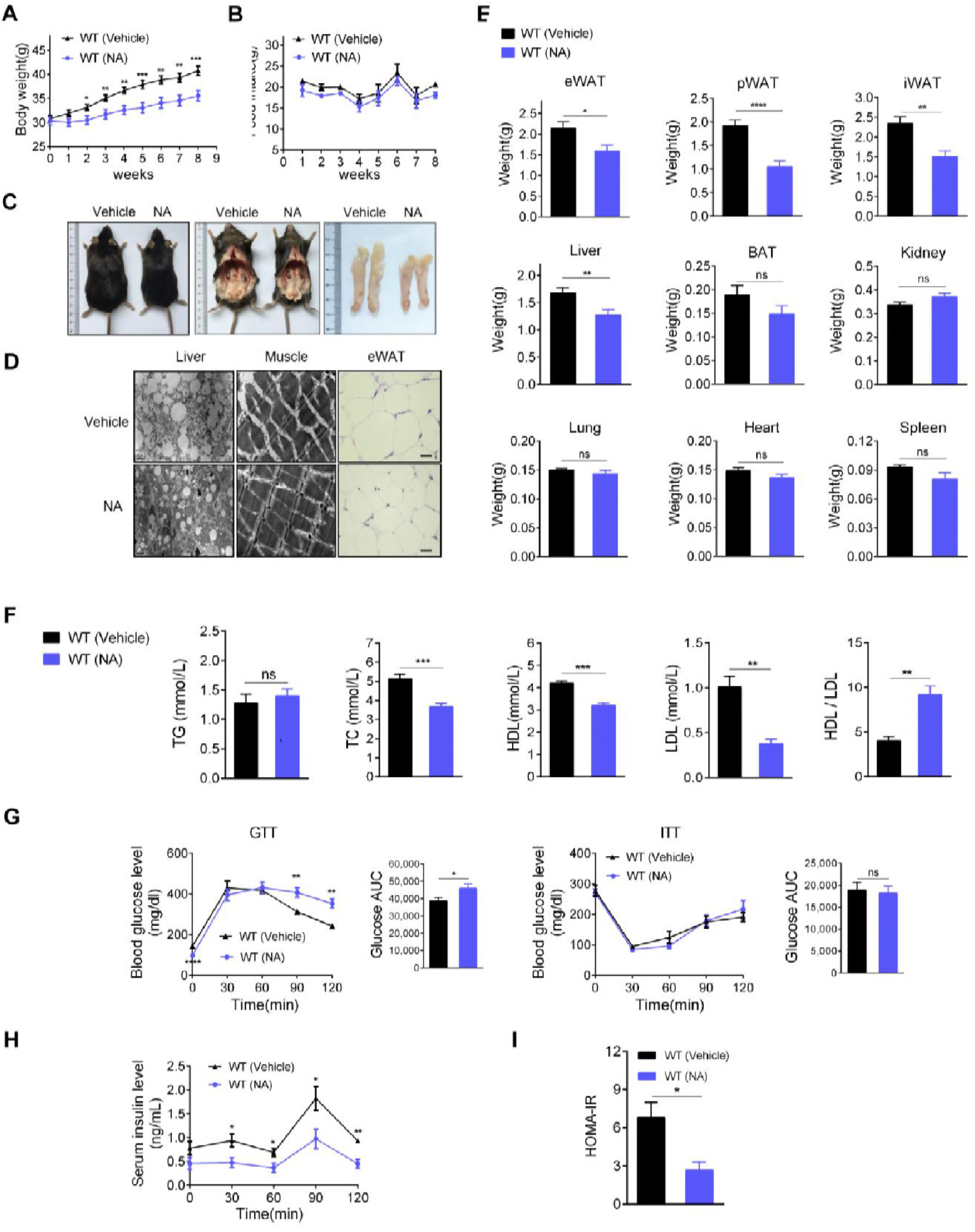
Niacin exerts inhibitory effects on HFD-induced obesity. A,B Body weight (A) and food intake (B) were recorded weekly (n=16-17). C Representative images of fat mass were shown. D Representative SEM images in liver (left panels; scale bars, as shown in figure) and muscle i (middle panels; scale bars, as shown in figure), and H<E staining of epididymal white adipose tissue (eWAT) (right panels; scale bars, 100 μm). ì E Fat pads and organ weights were recorded after mice anesthetized (n=16-17). i F Serum lipid parameters, including total triglyceride (TG), total cholesterol (TC), high density i lipoprotein (HDL), and low density lipoprotein (LDL) were measured using biochemical testing kits (n=7-8). G Glucose tolerance test (GTT) and insulin tolerance test (ITT) results and the corresponding; AUC for the blood glucose levels (n=5-6). H Serum insulin level during GTT was measured by ELISA (n=5). I HOMA-IR index was calculated as stated in the “Method” section (n=5). i Data information: All the data are presented as means ±SEM. In (A, B, E-I), data were analyzed by unpaired two-tailed student’s t-test. \**P* < 0.05; \*\**P* <0.01; \*\*\**P* < 0.001; \*\*\*\**P* < 0.0001.

Niacin has profound and unique beneficial effects on blood lipid (Carlson, 2005). We then examined effects of niacin challenge on metabolic parameters in serum of experimental mice. As shown in Fig 2, challenge with niacin led to a significant reduction in serum total cholesterol (TC), low-density lipoprotein (LDL) and high-density lipoprotein (HDL) with a characteristic of high ratio of HDL/LDL in mice fed on HFD, but triglyceride (TG) remained unchanged in respect to vehicle-treated mice. The effect of niacin on TC, LDL, HDL and ratio of HDL/LDL was not observed in Gpr109a^−/−^ mice and mice challenged with CHBA, a specific small molecule agonist for GPR81 (Figs 2F and EV2A). We also performed glucose tolerance test (GTT) and insulin tolerance test (ITT) and documented that niacin treated mice were less glucose tolerant, but exhibited insulin tolerant comparable to control mice (Fig 2G). No changes on GTT and ITT were detected in niacin-treated Gpr109a^−/−^ mice and CHBA-challenged wild-type mice (Fig EV2B). Quantitative analysis showed that niacin treatment results in a significant reduction in blood insulin (Fig 2H), however, challenge of HFD-fed mice with niacin exhibited a markedly decrease in HOMA-IR values compared to vehicle-treated mice (Fig 2I), suggesting the beneficial role of niacin in insulin sensitivity.

### Lack of GPR109A prevents anti-obesity actions of niacin

To further validate the role of GPR109A in the anti-obesity action of niacin, we challenged HFD-fed Gpr109a^−/−^ male mice with niacin. As indicated in Fig 3, Challenge of HFD-fed Gpr109a^−/−^ mice with niacin exhibited similar weight gain, adipose tissue weight, organ weight, and food intake as respect to vehicle-treated Gpr109a^−/−^ mice (Fig 3A-C, and E). Histological observation also showed no difference in lipid accumulation between niacin-treated and vehicle-treated Gpr109a^−/−^ mice (Fig 3D). These results suggest that niacin exerts its regulatory effects on lipid metabolism through GPR109A.

**Figure 3.**
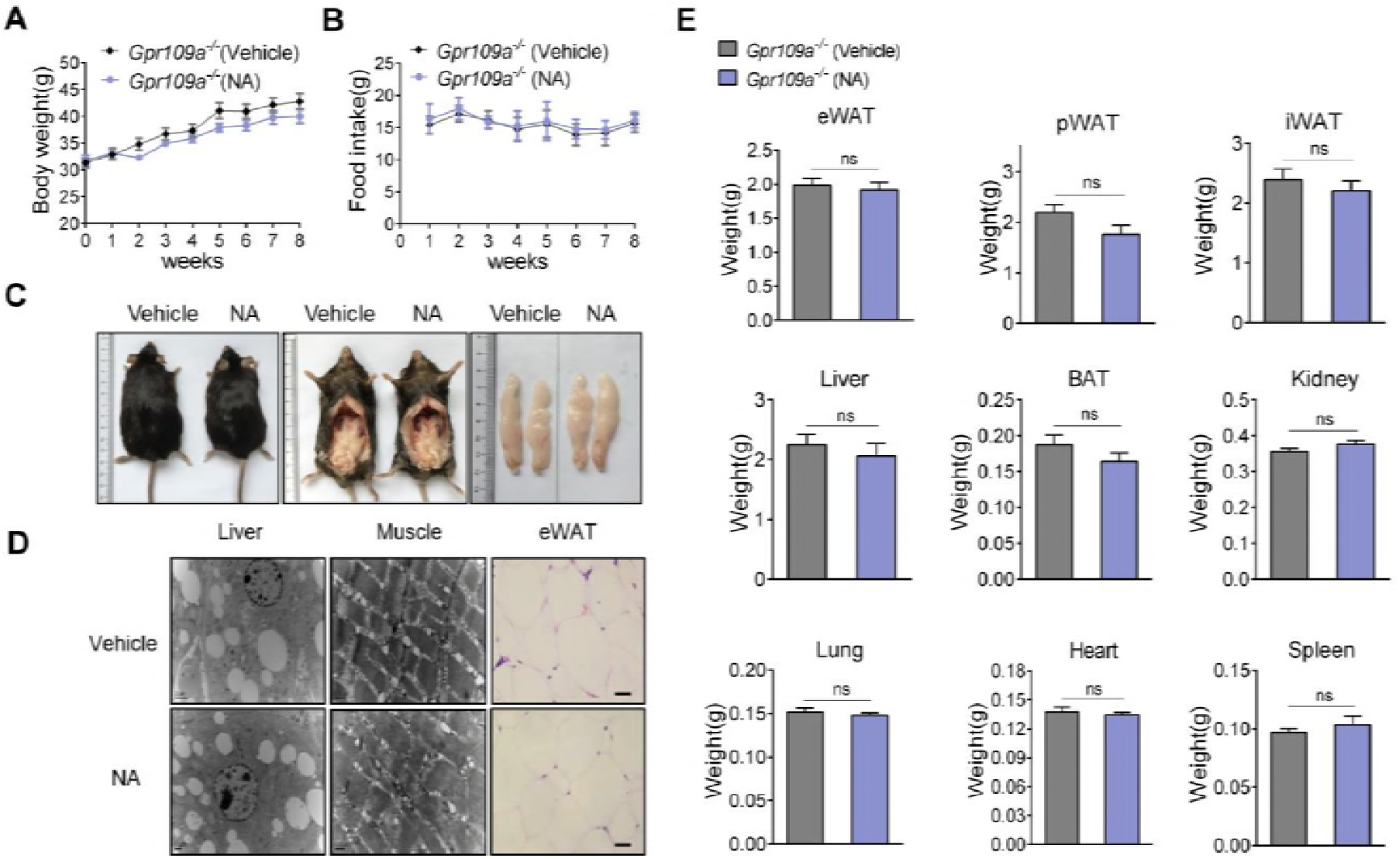
Lack of GPR109A prevents anti-obesity actions of niacin. A,B Body weight (A) and food intake (B) were recorded weekly (n=10-11). C Representative images of fat mass were shown. D Representative SEM images in liver (left panels; scale bars, as shown in figure) and muscle (middle panels; scale bars, as shown in figure), and H&E staining of epididymal white adipose tissue (eWAT) (right panels; scale bars, 100 μm). E Fat pad and organ weights were recorded after mice anesthetized (n=10-11). Data information: All the data are presented as means ±SEM. In (A, B, E), data were analyzed by unpaired two-tailed student’s t-test. ns, not significant.

### Niacin directly inhibits hepatocyte lipogenesis through GPR109A

Our data clearly showed that upon activation by niacin, GPR109A exerts its inhibitory effects on lipid accumulation in adipose tissue, liver and muscle when fed on a HFD. Moreover, oil red O staining and quantitative analysis revealed that HFD-fed mice displayed a significant increase in hepatic triglyceride content, whereas niacin supplement led to a marked reduction in triglyceride content in liver (Fig 4A and B), in contrast, no difference in TC and FFA levels was observed (Fig 4B). In addition, the decrease in the hepatic insulin level was observed in mice with niacin challenge (Fig 4C), coincident with the result in serum. However, animals remain unaltered in food intake. We then investigate whether or not niacin plays a role in the regulation of lipogenesis via GPR109A. We first used qRT-PCR to quantitatively analyze the GPR109A expression in mouse liver.

**Figure 4.**
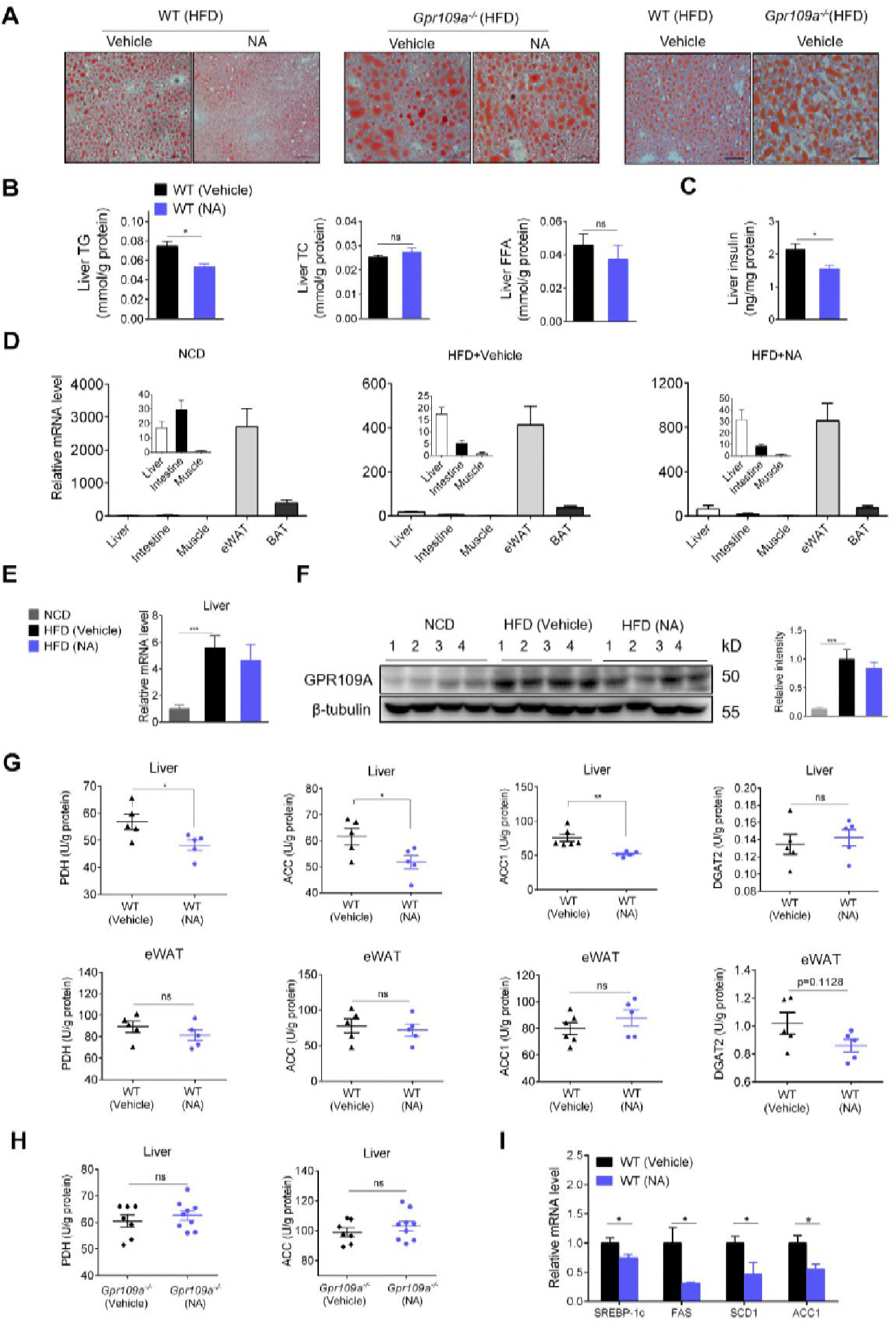
Niacin directly inhibits hepatocyte lipogenesis through GPR109A. A Representative images of Oil Red O staining of livers. Scale bars, 100 μm. B Liver lipid content, including the levels of TG, TC, and FFA, were measured (n=5-6). C Insulin levels were measured by ELISA (n=5). D GPR109A gene expression was measured by qRT-PCR in liver, intestine, muscle, eWAT, and BAT of the mice fed normal chow diet (NCD), high-fat diet (HFD), and HFD with 50 mM niacin (NA) respectively. Gene expression in muscle served as the statistical controls (n=8-9). E Expression of GPR109A was measured in liver isolated from the WT mice fed NCD, HFD with or without NA by quantitative analyses (n=8-9). F The relative gene expression of GPR109A in liver was calculated by western blot (left panel) and quantitative analyses (right panel). The β-tubulin was used as a loading control. G Comparing the activities of enzymes involving in lipogenesis (PDH, ACC, ACC1, and DGAT2) of WT mice fed HFD with or without NA in liver (upper panels) and eWAT (lower panels) (n=5). H The activities of PDH and ACC in liver were measured in Gpr109a^−/−^ mice fed HFD (n=7,9). I The mRNA expression analysis of genes involved in de novo lipogenesis in liver, and the data of WT (Vehicle) group served as the statistical controls (n=8). Data information: All the data are presented as means ±SEM. In (B, C, E-I), data were analyzed by unpaired two-tailed student’s t-test. \**P* < 0.05, \*\**P* <0.01, \*\*\**P* < 0.001.

It has been reported that GPR109A is highly expressed in adipocytes and activated immune cells (Carlson, 2005; Soga et al, 2003; Tunaru et al, 2003; Wise et al, 2003). However, our data clearly showed that when fed with HFD, mice exhibited a significant increase in lipid accumulation in liver, whereas niacin supplement resulted in a significant reduction of lipid accumulation in liver. To confirm the role of GPR109A in liver, we examined the expression of GPR109A in different tissues and organs. High-level expression of GPR109A mRNA was detected in epididymal WAT and BAT (Fig 4D). Interestingly, although relatively lower basal expression of GPR109A was detected, upon challenge with HFD, GPR109A mRNA was markedly upregulated in liver, while niacin stimulation showed no effect on GPR109A expression in different tissues and organs (Fig 4E and EV3A). In contrast, GPR109A was completely absent in liver from Gpr109a^−/−^ mice (Fig EV3B and C). GPR109A expression in liver was also confirmed at protein level by western blot (Fig 4F). However, our data on GPR109A expression in mouse liver could not discriminate between the hepatocytes and Kupffer cells. Therefore, to confirm the expression of GPR109A in hepatocytes, we further analyzed the expression and function of GPR109A in HepG2 cells, a human liver carcinoma cell line endogenously expressing GPR109A (Fig EV4A). As anticipated, niacin (300 μM) induced remarkable phosphorylation of ERK1/2 in a time-dependent manner with a maximal activation at 5 min (Fig 4B). In addition, niacin (100 μM) induced Ca^2+^ mobilization in HepG2 cells (Fig EV4C). These data suggest that GPR109A is functionally expressed in hepatocytes.

To confirm the role of GPR109A in the regulation of de novo lipogenesis in liver, we used the ELISA to quantitatively analyze the activities of key enzymes involved in fatty acid biosynthesis, acetyl CoA carboxylase (ACC), acetyl CoA carboxylase-1 (ACC-1), and pyruvate dehydrogenase (PDH) (Tong, 2005). As shown in Fig 4G, in wild-type mice fed with HFD, treatment with niacin resulted in a significant decrease in the enzyme activities of ACC, ACC1, and PDH in liver, while no change in WAT was observed. However, in Gpr109a^−/−^ mice, no difference in the enzyme activities of PDH and ACC in liver followed treatment with niacin was detected (Fig 4H). Our data also showed that niacin supplement led to no statistically significant changes in the activity of diglyceride acyltransferase 2 (DGAT2) (Fig 4G), which directly catalyzes the formation of triglycerides from diacylglycerol and Acyl-CoA in fat, liver and skin cells (Oelkers et al, 1998). In addition, these data were further confirmed by quantitative determination of mRNA transcripts of sterol regulatory element-binding protein-1c (SREBP-1c), a key transcriptional activator of lipogenic genes, and its downstream target genes, fatty acid synthase (FAS), stearoyl-CoA desaturases-1 (SCD-1) and acetyl CoA carboxylase-1 (ACC-1), by real time RT-PCR. As revealed in Fig 4I, administration of niacin resulted in a significant decrease in mRNA levels of SREBP-1c and its target genes, FAS, SCD-1 and ACC-1, in liver compared to the vehicle-treated mice.

### AMPK and ERK1/2 are involved in the GPR109A-mediated inhibition of hepatocyte lipogenesis

Next, to elucidate the mechanism(s) involved in niacin-initiated inhibition of hepatocyte lipogenesis through GPR109A, western blot analysis was performed to determine the activation of AMP kinase (AMPK), extracellular signal-regulated kinase (ERK) and protein kinase B (Akt). As shown in Fig 5A, niacin-treated wild-type mice exhibited a markedly enhancement in phosphorylation of AMPK, ERK1/2 and Akt in liver, while only phosphorylation of AMPK in muscle, but not in eWAT. However, niacin-treated Gpr109a^−/−^ mice had no detectable differences in the activation of AMPK, ERK1/2 and Akt in liver (Fig 5B). The cultured HepG2 cells, a human liver carcinoma cell line endogenously expressing GPR109A (Fig EV4A), were used to further examine the GPR109A-mediated intracellular signaling. As shown in Fig 5C, as anticipated, niacin (300μM) induced remarkable phosphorylation of AMPK, ERK1/2, and Akt in a time-dependent manner with a maximal activation at 5 min. Accordingly, phosphorylation and inactivation of ACC, a downstream target of AMPK, was observed (Fig 5C). Consistently, siRNA-mediated knockdown of GPR109A expression led to the significant impairment of niacin-induced ERK1/2, AMPK, and ACC phosphorylation and Ca^2+^ mobilization in HepG2 cells (Figs 5D and EV4D-F). Taken together, these data suggest that niacin exerts its inhibitory effect on ACC through GPR109A-mediated activation of AMPK.

**Figure 5.**
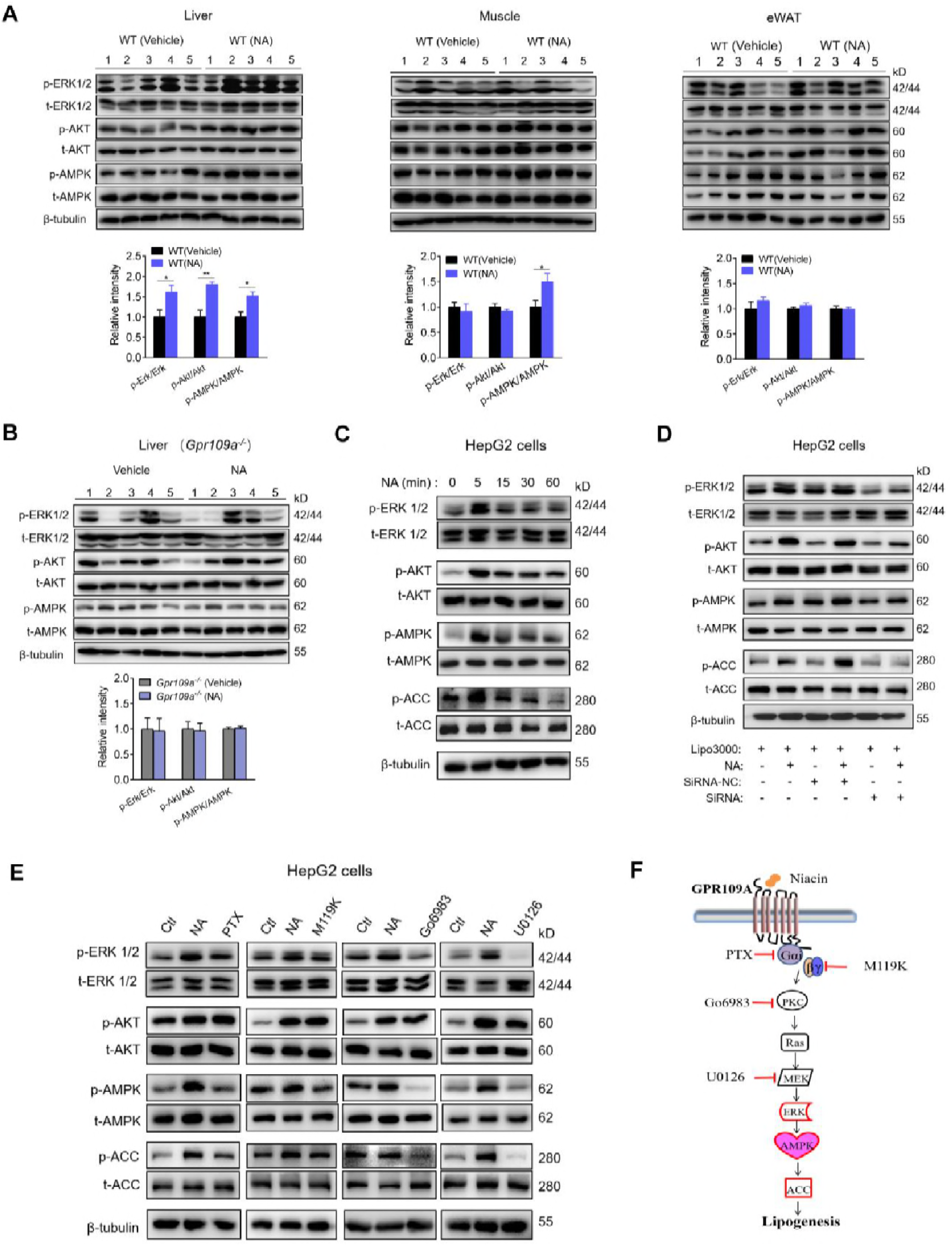
AMPK and ERK1/2 are involved in the GPR109A-mediated inhibition of hepatocyte lipogenesis. A Phosphorylated of ERK1/2, AKT and AMPK were measured in the liver, muscle, and eWAT from the WT mice by western blot and quantitative analyses (n=5). B Phosphorylated of ERK1/2, AKT and AMPK were measured in the liver from Gpr109a^−/−^ mice. (n=5). C Activation of ERK1/2, AKT, AMPK and ACC in serum-starved HepG2 cells stimulated with niacin (300 μM) for the indicated time periods. D HepG2 cells were transfected with GPR109A siRNA for 36 h and stimulated with niacin (300 μM) for 5 min. Phosphorylation of ERK1/2, AKT, AMPK and ACC was accessed. E Activation of ERK1/2, AKT, AMPK, and ACC in serum-starved HepG2 cells pretreated with PTX (100 ng/mL) for 16 hr or M119K (10 μM), Go6983 (10 μM), or U0126 (1 μM) for 1h followed by a challenge with niacin (300 μM) for 5 min was accessed by western blot. And β-tubulin was used as a loading control. F Schematic model of suppression of ACC via GPR109A. Data information: All the data are presented as means ±SEM. In (C,D,E), All data of cells shown are representative of at least three independent experiments. In (A,B), data were analyzed by unpaired two-tailed student’s t-test. \**P* < 0.05, \*\**P* <0.01.

We used western blot analysis combined with various inhibitors of G proteins and kinases to explore the role of ERK1/2 and/or AKT in the activation of AMPK. Our results showed that inhibitors of PTX for Gαi, M119K for Gβγ, Go6983 for PKC, and U0126 for MEK exhibiting inhibitory effects on ERK1/2 phosphorylation exerted the same inhibitory activities on AMPK (Fig 5E). Collectively, it is likely that niacin exhibits inhibitory effect on hepatic lipid synthesis through activation of AMPK via Gβγ-PKC-dependent ERK1/2 signaling pathway, as the schematic model shown (Fig 5F).

### Niacin treatment triggers BAT thermogenesis via GPR109A

Obesity is an abnormal accumulation of lipid in adipose tissue caused by an imbalance between energy intake and expenditure. In HFD-induced obesity mouse model, niacin supplement led to a significant reduction in body weight, but no change in daily food intake, indicating that niacin is more likely to promote energy expenditure. To validate this hypothesis, we first monitored the core body temperature by measuring the anal temperature of mice fed an HFD and treated with vehicle or niacin upon acute cold exposure. GPR109A-deficient mice showed a continuous decline in body temperature and reached 32°C after acute cold exposure at 4°C for 6 hours, while wild-type mice were able to maintain their body temperature at 33.5°C (Fig 6A). Moreover, wild-type mice fed an HFD and treated with niacin revealed relatively but significantly higher body temperature than that of vehicle-treated mice, however, no significant difference was observed between niacin- and vehicle-treated group in GPR109A-deficient mice in response to acute cold exposure (Fig 6A) and in wild-type mice without acute cold exposure (Fig 6B). These data collectively confirm cold tolerance in niacin-challenged wild-type mice.

**Figure 6.**
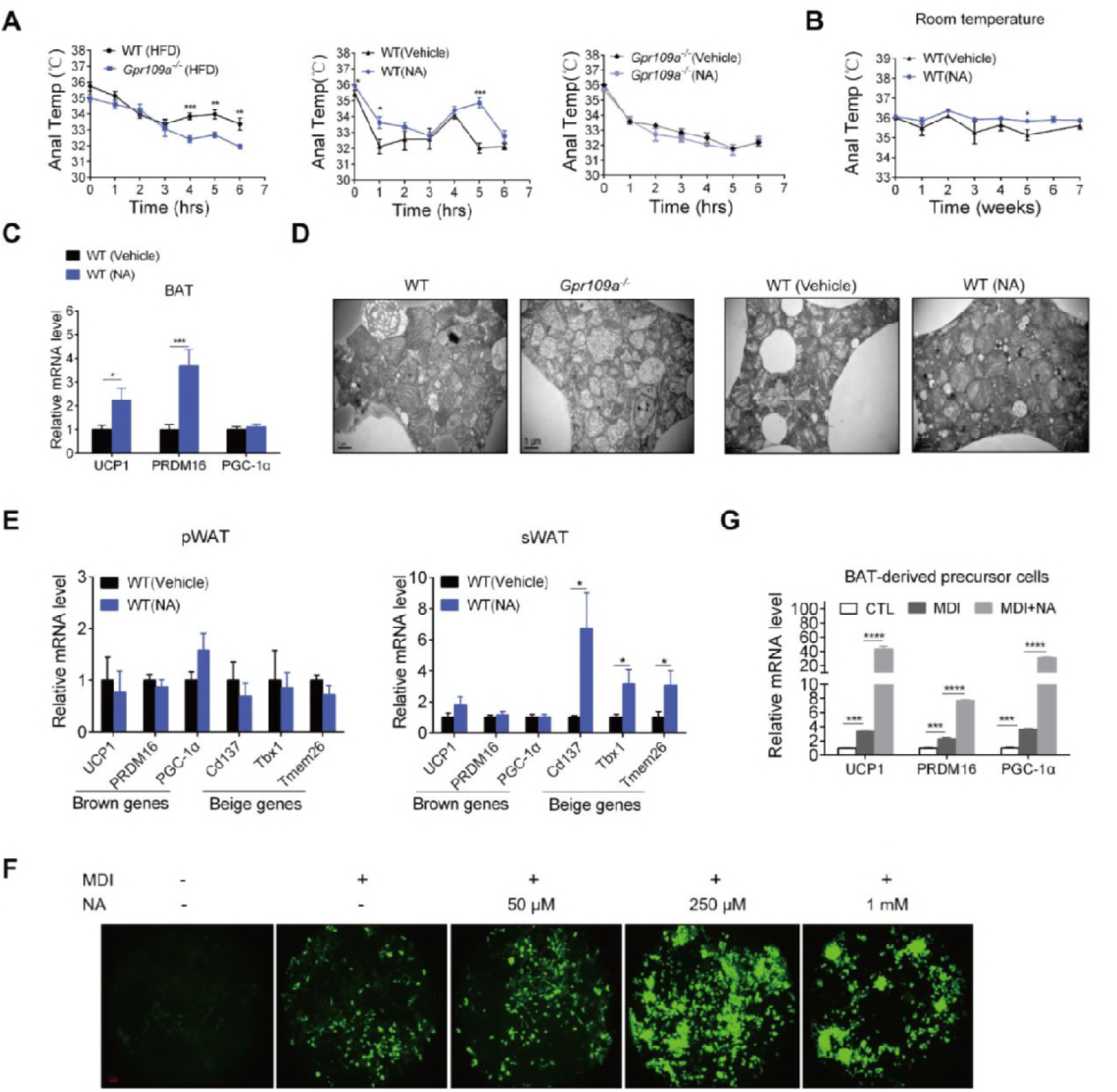
Niacin treatment triggers BAT thermogenesis via GPR109A. A Mice were exposed to 4°C for 6 hours. Monitor the anal temperature every hour (n=5-6). B Monitor the anal temperature weekly under the room temperature (n=5). C The mRNA expression of UCP1, PGC-1α and PRDM16 in brown adipose tissue (BAT) from the WT mice (n=11-15). D Electron micrographs of the BAT at the end of the experiment. Scale bar: 1μm. E The mRNA expression of brown or beige genes (UCP1, PRDM16, PGC-1α, Cd137, Tbx1, and Tmem26) in eWAT and sWAT (n=10-11). F Brown fat precursor cells were differentiated in MDI with or without niacin for 8 days. Visible lipid accumulations were observed by BODIPY 493/503 staining. Scale bars, 100 μm. G The mRNA expression of UCP1, PRDM16, and PGC-1α in MDI-induced brown fat precursor cells with the indicated concentration of niacin for 8 days. Data information: All the data are presented as means ±SEM. In (F,G), All data of cells shown are representative of at least three independent experiments. In (A-C, E, G), data were analyzed by unpaired two-tailed student’s t-test. \**P* < 0.05, \*\**P* <0.01, \*\*\**P* < 0.001, \*\*\*\**P* < 0.0001.

Brown adipose tissue (BAT) is the major site essential to generate heat (thermogenesis) for the maintenance of body temperature. Therefore, we next examined the expression of genes involved in BAT thermogenesis real time RT-PCR analysis. As shown in Fig 6C, the mRNA expression of uncoupling protein 1 (UCP1), a critical player in allowing electrons to be released as heat rather than stored, and positive regulatory domain 16 (PRDM16), a well-known BAT transcriptional regulator, were significantly up-regulated in HFD-fed wild-type mice in response to challenge of niacin. This is further confirmed by observation of the ultrastructure of BAT mitochondria. Transmission electron microscopy demonstrated that BAT from Gpr109a^−/−^ mice exhibited significant reduction in number and size of mitochondria with aberrant cristae compared to wild-type mice. Niacin challenge caused a remarkable increase in number and size of mitochondria with improved cristae organization in HFD-fed wild-type mice (Fig 6D). Additionally, we also observed that niacin showed the potential to rapidly elevate the expression of beige-specific genes including Cd137, Tbx1, and Tmem26 in sWAT (Fig 6E). These data collectively suggest that niacin stimulates BAT energy expenditure through GPR109A in HFD-fed mice.

Additionally, we also evaluated the role of niacin/GPR109A signaling in the preadipocyte differentiation. Niacin showed stimulatory effect on the differentiation-induction of brown fat precursor cells with a maximal activity at 250 μM (Fig 6F). Further real-time RT-PCR analysis revealed that niacin treatment significantly up-regulated the transcript levels of UCP1, PGC-1a and PRDM16 (Fig 6G).

### Niacin stimulation decreases fatty acid absorption though GPR109A

Recent studies have demonstrated that GPR109A is expressed in colonic and intestinal epithelial cells and mediates the local control of intestinal glucose absorption (Wong et al, 2015) and the tumor-suppressive effects of the bacterial fermentation product butyrate in colon (Thangaraju et al, 2009). Accordingly, we further examined the role of GPR109A expressing in colonic and intestinal epithelial cells in the control of intestinal fatty acid absorption. We collected feces from mice housed individually from 5 consecutive days and measured fecal lipid contents. As indicated in Fig 7, the weights of feces normalized to body weight in the niacin-challenged mice when fed on HFD compared to the control mice (Fig 7A). Further analysis showed that total lipid, TG, TC and FFA in feces were significantly increased when HFD-fed mice challenged by niacin compared to the control mice. However, total lipid in feces was reduced in HFD-fed Gpr109a^−/−^ mice treated with niacin, no difference in TG, TC and FFA was observed between niacin-challenged mice and control mice (Fig 7B). This was confirmed by RT-PCR quantitative analysis of three key proteins (FATP4, I-FABP, and NCP1L1) involved in the control of fatty acid and cholesterol absorption. The mRNA levels of the three genes were obviously decreased in the wild-type NA group, but not in the Gpr109a^−/−^ mice (Fig 7C). These findings suggest that niacin is likely to exhibit the potential to inhibit the dietary fat absorption through GPR109A, thereby causing the decrease in fat deposit and body weight.

**Figure 7.**
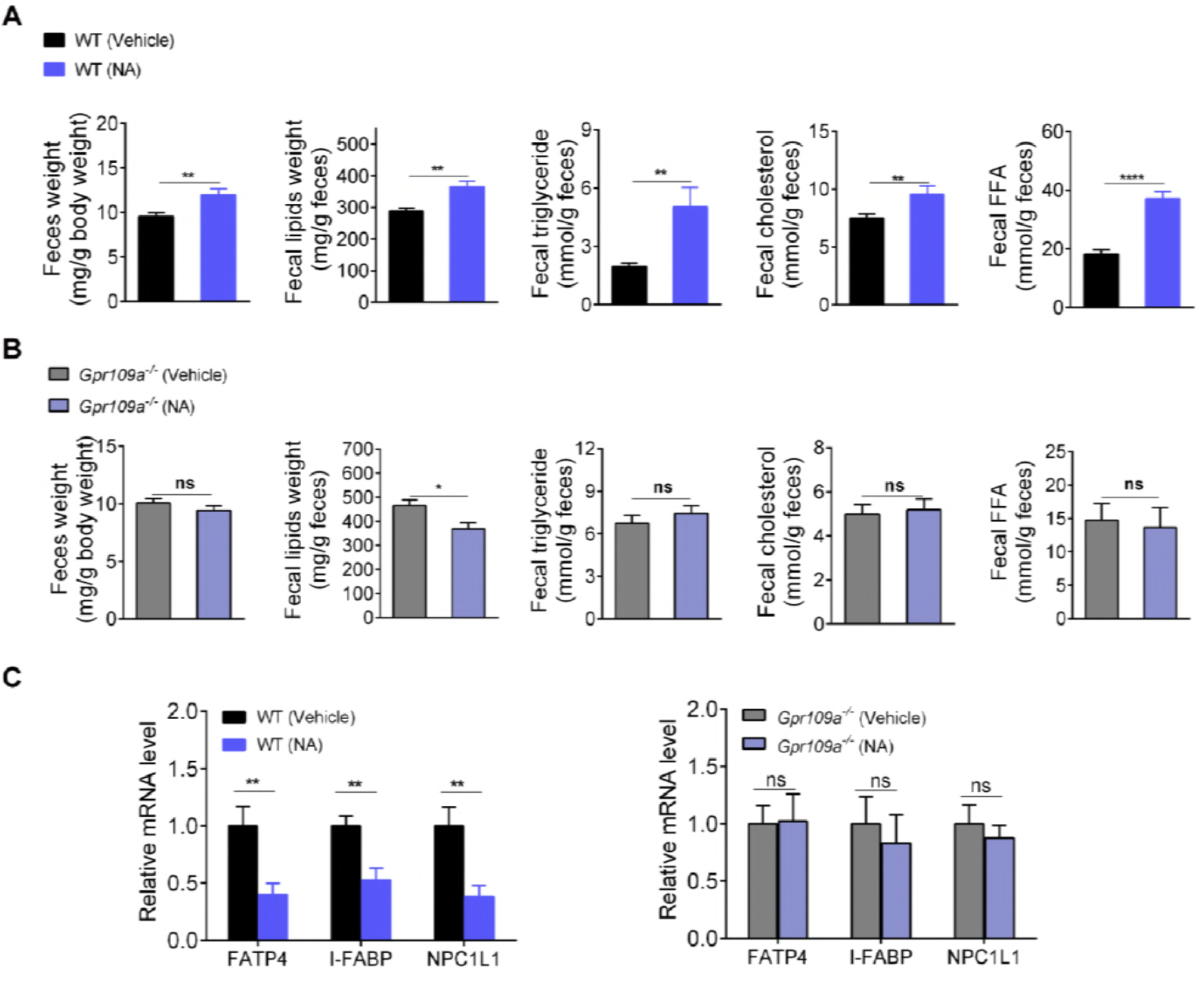
Niacin stimulation decreases fatty acid absorption though GPR109A. **A** Fecal mass were normalized to body weight (mg/g). Fecal lipids were extracted and the fecal lipids concentration was expressed as milligram of lipids per gram of feces weight. Total triglyceride, total cholesterol, and free fatty acid (FFA) were measured in WT mice (n=7,9). **B** Feces lipid indicators were measured in Gpr109a^−/−^ mice (n=6). **C** Total mRNA isolated from intestines and examined the mRNA expression of indicated genes involved in fatty acid absorption (fatty acid transport protein 4 (FATP4) and intestinal-type fatty acid-binding protein (I-FABP)) and cholesterol absorption (Niemann-Pick C1-Like 1 (NPC1L1)) by qRT-PCR. Data information: All the data are presented as means ±SEM. In (F,G), All data of cells shown are representative of at least three independent experiments. Data were analyzed by unpaired two-tailed student’s t-test. \**P* < 0.05, \*\**P* <0.01.

## Discussion

Since the identification of GPR109A, a Gi/o-coupled receptor, as the molecular target for niacin (Digby et al, 2012; Ganji et al, 2009; Lukasova et al, 2011b), most studies have focused on its pronounced hypolipidemic and cardioprotective action. Recent investigations have also shed light on anti-inflammatory and anti-oxidative effects (Buhler et al, 2018; Digby et al, 2012; Gautam et al, 2018; Graff et al, 2016; Singh et al, 2014). However, little attention has been paid to the potential of this receptor to favorably modulate lipogenesis and energy homeostasis. In the present study, we explored physiological roles of niacin/GPR109A in the regulation of lipid metabolism and energy homeostasis in a high-fat diet-induced murine model of obesity combined with GPR109A deficient mice. The most important observation is that niacin supplementation prevented the progression of dietary obesity in HFD-fed wild mice, but not in HFD-fed GPR109A deficient mice. Mechanistically, we demonstrated that niacin-mediated reduction in body weight and fat mass was caused by fine-tuning of hepatic de novo lipogenesis, BAT thermogenesis and intestinal fatty acid absorption through GPR109A.

During the breeding process of GPR109A knockout mice introduced from Dr. Offermanns’ lab, we observed that the deletion of GPR109A resulted in obvious increase in body weight and fat mass compared to wild-type mice. In contrast, lactate receptor GPR81-deficient mice gain significantly less body weight than their wild-type under HFD-feeding conditions, although Gi/o-coupled GPR81 exerts same activity in the inhibition of lipolysis in white adipose tissue (Ahmed et al, 2010). This prompted us to initiate this study to evaluate the exact role of GPR109A in the regulation of energy homeostasis. A recent study indicated that niacin treatment remarkably reduced body weight in the obese mice (Chen et al, 2015b). Previous investigations have demonstrated that the ketone body β-hydroxybutyrate (βOHB), as an important metabolic intermediate and also a gut bacterial fermentation-yielded product, which is the only known endogenous ligand for GPR109A (Taggart et al, 2005), causes reductions of body weight and fat content in rats and mice (Davis et al, 1981; Gao et al, 2009). Using combination of GPR109A deficient mice with HFD-induced model of obese, we provided further evidence that niacin treatment led to a significant reduction in body weight and fat mass through GPR109A when fed on normal chow diet or HFD. These findings suggest that GPR109A has the potential to control energy homeostasis, preventing weight gain.

Our results clearly revealed that niacin challenge remarkably reduced liver weight in wild-type mice fed on HFD, but not in GPR109A-deficient mice. Ultrastructural examination validated that GPR109A-deficient mice had a striking visible increase in amount and size of lipid droplets in liver compared with the wild-type group, and niacin treatment led to a significant reduction in amount and size of lipid droplets in liver when fed on HFD. These findings suggest that GPR109A play an important role in the regulation of de novo lipogenesis in liver. The liver has long been recognized as the major site of de novo lipogenesis, responsible for the conversion of excess dietary carbohydrate into triglycerides. De novo lipogenesis turnover accounts for about 20% of lipid turnover within adipose tissue (Strawford et al, 2004), however, De novo lipogenesis in non-adipose tissues leads to ectopic lipid accumulation responsible for metabolic stress and insulin resistance (Solinas et al, 2015). Although expression of GPR109A in liver remains controversial, the abundance of GPR109A mRNA was detected higher in liver in lean-type than in fat-type cows (Mielenz et al, 2013). GPR109A has been found to be expressed in murine liver but the basal expression is low (Li et al, 2010). In the current study, we provided evidence to confirm that GPR109A mRNA and protein were detected in murine liver at relatively lower level, however, mice fed on HFD exhibited significantly enhanced expression of GPR109A in the liver. In addition, the functional expression of GPR109A in HepG2 cells was also detected. Further biochemical analysis of key enzymes involved in lipogenesis demonstrated that niacin exhibited inhibitory effects on PDH, the gatekeeping enzyme of pyruvate flux into acetyl-CoA, and ACC1, the key regulatory step of fatty acid synthesis (Katsurada et al, 1990; Ruderman et al, 2003). These findings were verified by real-time qRT-PCR for genes encoding SREBP-1c, a key transcriptional activator of lipogenic genes, and lipogenic enzymes, FAS, SCD1 and ACC1 (Rui, 2014; Shi & Burn, 2004; Titchenell et al, 2015). Niacin has been shown to directly inhibit the activity of diglycerol acyltransferase 2 (DGAT2), the key enzyme that catalyzes the final step in triglyceride synthesis in liver (Ganji et al, 2004; Kamanna et al, 2013). However, our results strongly suggest that niacin exerts its inhibitory effects on hepatic lipogenesis through GPR109A-mediated signaling.

Moreover, mechanistically, we demonstrated that Akt, ERK1/2 and AMPK were involved in niacin/GPR109A-mediated inhibition of hepatic lipogenesis. AMP-activated protein kinase (AMPK), a serine/threonine kinase, has been recognized as an intracellular energy sensor for maintaining energy homeostasis (Shackelford & Shaw, 2009). AMPK activation is associated with the hepatic inhibition of de novo lipogenesis by modulating transcription factor SREBP-1c and lipogenic key enzyme ACC1 (Li et al, 2011b; Zhou et al, 2001). To gain further insight into the mechanism underlying the GPR109A-mediated inhibition of hepatic lipogenesis and lipid accumulation, we examined the GPR109A-midiated signaling pathways responsible for the niacin-triggered AMPK activation. We demonstrated that MEK inhibitor U0126 significantly suppressed ERK1/2 phosphorylation and AMPK activation, suggesting that the activation of ERK1/2 is likely upstream of AMPK activation. Additionally, treatment with Gi/o inhibitor PTX, Gβγ inhibitor M119K and PKC inhibitor Go6983, respectively, resulted in a remarkable decrease in both ERK1/2 and AMPK phosphorylation. Collectively, these findings suggest that niacin causes inhibition of hepatic lipogenesis via GPR109A-dependent Gβγ/PKC/ERK1/2/AMPK signaling pathway.

It is noteworthy that mice challenged with niacin exhibited a remarkable reduction in body weight and fat mass without affecting food intake, consistent with previous observation that βOHB supplementation elicited body weight loss but without a decrease in food intake (Davis et al, 1981; Gao et al, 2009). It suggests that energy expenditure is likely to contribute to niacin-induced body weight loss. Results derived from cold response experiments showed that thermogenic function is enhanced in wild-type mice challenged with niacin, but not in GPR109A knockout mice. Real time RT-PCR analysis demonstrated that treatment with niacin increased the expression of thermogenic genes UCP1, a thermogenic protein exclusively expressed in brown/beige adipocytes (Cannon & Nedergaard, 2004; Gesta et al, 2007), PGC-1α, a master regulator of mitochondrial biogenesis (Cohen et al, 2014), and PRDM16, a well-known BAT transcriptional regulator (Harms & Seale, 2013) in mice fed on HFD, and in MDI-induced brown fat precursor cells. Interestingly, we found that mitochondria in BAT of GPR109A knockout mice exhibited smaller in size with abnormal cristae structures compared to wild-type mice, and niacin challenge showed significant improvement in mitochondrial size and cristae structure of mice fed on HFD. Additionally, niacin stimulated a beige fat thermogenic program of the sWAT. These data strongly suggest that niacin treatment triggers BAT and/or browning WAT thermogenic activity in mice through GPR109A.

Importantly, GPR109A has been shown to functionally express in colonic and intestinal epithelial cells (Thangaraju et al, 2009; Wong et al, 2015). Niacin has been found to locally control glucose uptake at the level of the small intestine though GPR109A (Wong et al, 2015). This prompts us to examine whether or not GPR109A is involved in the control of uptake of sterols and fatty acids in the level of the small intestine. To our surprise, niacin supplementation caused a significant increase of total lipids, triglycerides, cholesterols, and free fatty acids (FFAs) in the feces of wild-type mice fed on HFD, but no change was observed in HFD-fed GPR109A-deficient mice. Further gene expression analysis showed that genes encoding FATP4, I-FABP and NPC1L1 exhibited a significant reduction in expression in the intestinal tissues. It is established that polytopic transmembrane protein NPC1L1 is essential for intestinal absorption of cholesterol (Altmann et al, 2004), whereas proteins FATP4 and I-FABP play a role in dietary fat absorption and fatty acid transport, respectively (Besnard et al, 2002; Hall et al, 2005). Taken together, these findings suggest that in addition to the control of glucose uptake, GPR109A expressed in the small intestine is likely involved in the local regulation of intestinal absorption of triglycerides, cholesterols, and free fatty acids.

Consistent with previous studies (Chang et al, 2006; Chen et al, 2015a; Garg & Grundy, 1990; Goldberg & Jacobson, 2008), we also observed that the HFD-induced obese mice treated with niacin exhibited an obvious reduction in serum insulin concentration and glucose tolerance compared to the vehicle-treated mice. However, our data demonstrated that treatment with niacin led to a significant improvement in insulin resistance in HFD-fed obese mice (HOMA-IR). GPR109A has been shown to be functionally expressed in both human and murine islet beta-cells (Li et al, 2011a; Wang et al, 2016). Chen et al. have shown that niacin exerts its inhibitory actions on β-cell function via GPR109A-mediated signaling, but without affecting islet architecture (Chen et al, 2015a). Nevertheless, a recent study demonstrated that GPR109A expression was down-regulated dramatically in islet β-cells of type 2 diabetic patients as well as diabetic db/db mice, suggesting that the important role of GPR109A signaling in the prevention of diabetes (Wang et al, 2016). Moreover, an early study observed that a short-term niacin analogue administration resulted in an improved glucose tolerance in prediabetic or diabetic patients (Fulcher et al, 1992). In addition, our data clearly demonstrated that niacin challenge caused a significant reduction in hepatic de novo lipogenesis and ectopic lipid accumulation in liver, adipose tissue and muscle of mice fed on HFD, leading to a beneficial effect on insulin resistance. The effects of niacin on glucose homeostasis remain controversial, and the exact roles of GPR109A in the regulation of insulin resistance now deserve thorough evaluation.

In conclusion, the data presented in this report have demonstrated a novel role of niacin in the regulation of energy homeostasis by fine-tuning of hepatic de novo lipogenesis, BAT/beige thermogenesis and intestinal fat absorption via GPR109A-mediated signaling. Our findings provide a critical clue to the potential new therapeutic strategies that could treat obesity, diabetes and cardiovascular disease. As the debate continues about the favorable effects of niacin/GPR109A on the development and progression of atherosclerosis and its complications, it will be vital to further perform a more comprehensive assessment of niacin/GPR109A in the regulation of energy homeostasis in addition to anti-inflammation.

## Materials and Methods

### Cell culture and treatment

HepG2 cells (ATCC) were maintained in Dulbecco’s modified Eagle’s medium (DMEM) (Hyclone, USA) containing 10% heat-inactivated fetal bovine serum (FBS) (GIBCO, USA) with 1% penicillin-streptomycin solution. All cells were incubated at 37°C in a humidified chamber with 5% CO2. For the western blot analysis, the HepG2 cells were seeded in 24-well plates. For examining the intracellular signaling pathway, all cells were starved for 5h in serum-free medium to reduce the background before the agonist activation. And after stimulation with niacin (Sigma, 300μM), cells were lysed by RIPA buffer. If inhibitors were required, they were added to the cell before agonist stimulation. All samples were stored at −20°C until use.

### Small Interfering RNAs (SiRNAs) Transfection

Small-interfering RNA (siRNA) against GPR109A and a non-specific scrambled siRNA (siRNA-NC) were chemically synthesized by Dharmacon RNA Technologies (Lafayette). The sequences of siRNA were 5’ UAUGUGAGGCGUUGGGACU-3’ for GPR109A and 5’-UCAAAUAACCAUUCCAA GA-3’ for negative control. Cells at 60% confluence were transfected with siRNA using LipofectamineTM 3000 reagent (Invitrogen, USA), according to the manufacturer’s specifications. Media were replaced with DMEM containing 10% heat-inactivated FBS 6 h later. After the siRNA transfection, the cells were split for the indicated assay the following day.

### Brown fat precursor cells extraction and culture

Brown fat precursor cells were isolated from interscapular BAT of newborn wild-type mice as previously described (Cannon & Nedergaard, 2001). BAT was stripped from mice and transplanted into 6 cm cell culture dish with PBS immediately. Then cut the tissue into small pieces with elbow surgery. Disperse the broken tissue with 5mL isolation buffer containing Collagenase I in 37°C incubator for 30 min. After digestion, tissue remnants were removed by filtration through a 70μm nylon mesh (Corning, USA) and centrifuged at 200g for 5 min. Re-suspended with 5mL isolation buffer without collagenase and filtered through a 40μm nylon mesh (Corning, USA) and centrifuged at 200g for 5 min. Lastly, re-suspended cells with DMEM containing 10% FBS and inoculated into 6-well plate. When cells reached to 80%-90% density, began to induce differentiation.

### Brown fat precursor cells differentiation

For differentiation, two days after cells confluence (DAY 0), the medium was replaced with DMEM containing 10% FBS with inducers (MDI) (1.0μM dexamethasone, 10μg/mL insulin, and 0.5mM 3-isobutyl-1-methylxanthine) for 2 days. Two days after MDI (DAY 2), the medium was changed to DMEM supplemented with 10% FBS, 10μg/mL insulin. Two days later (DAY 4), cells were maintained in DMEM with 10% FBS for other 4-6 days. Full differentiation is usually achieved by DAY 8. Adipogenesis was monitored by morphological examination with Bodipy 493/503 (Molecular probes) staining.

### Animal studies

All mice were housed in groups of 4–6 per cage under constant temperature (23-25°C) with a 12 h light/12 h dark cycle. Animals were given ad libitum access to water and food. GPR109A heterozygous mice on C57BL/6 background were a generous gift from Dr. Offermanns’ lab (Bad Nauheim, Germany). Homozygous GPR109A knock out and their littermate wild-type mice were generated by matting heterozygous mice and genotyped using PCR. Male mice were used for all the experiments described in this study. And 4-weeks male C57BL/6 mice were purchased from the company of SLAC Laboratory Animal (Shanghai, China), and after a week of adaptation, mice were fed a HFD (60% kcal from fat) (D12492, Research Diets) for generating diet-induced obese (DIO) model and a normal chow diet (10% kcal, SLAC, Shanghai, China) in the control group. After 6 weeks on HFD, mice were divided randomly into two groups. One of group received niacin (50 mM) (Sigma-Aldrich, USA) dissolved in drinking water, and the other received vehicle (water) for 8-9 weeks as a control. Groups of age- and weight-matched animals were used in all experiments. Food consumption and body weight were recorded weekly. After the modeling period, mice were fasted overnight (12 h) and anesthetized with 3% isoflurane in a dedicated chamber, and the weights of major organ including fat pads and liver were recorded. Whole blood was collected and processed to isolate serum and tissues were flash frozen in liquid nitrogen and stored at −80°C for biochemical and histological analyses. All animal procedures were conducted in accordance with the Guide for the Care and Use of Laboratory Animals (United States National Institutes of Health). The protocol was approved by the research ethics committee.

### Body temperature in cold response

To test resistance to cold exposure, mice were individually caged and exposed to 4°C for 6 hours with free access to water. Anal temperature was monitored at the beginning and hourly after the start of cold exposure.

### Serum collection and parameters determination

Blood were collected form the orbital sinus, and samples were incubated at room temperature for 30 min to allow clotting. Serum samples were obtained after centrifuging at 3,500 rpm at 4°C for 20 min. Finally the supernatant (serum) was collected and stored at −80°C. Serum total triglyceride (TG), total cholesterol (TC), high density lipoprotein (HDL), and low density lipoprotein (LDL) were measured by using biochemical testing kits (Roche, Switzerland). Free fatty acids (FFA) were assayed by commercial kits (Nanjing Jiancheng, China). And plasma insulin levels were determined by a rat/mouse insulin enzyme-linked immunosorbent assay (ELISA) kit (Millipore Billerica, USA) according to the manufacturer’s instruction. Homeostasis model assessment of insulin resistance (HOMA-IR) was calculated using the following equation: HOMA-IR = (fasting plasma insulin (mIU/L) × fasting plasma glucose (mmol/L)/22.5.

### Fecal and liver lipids analysis

Fecal lipids were extracted as previously described (Folch et al, 1957). Feces were collected from mice housed individually from 5 consecutive days. Feces were dried for 1 h at 70°C, and one-hundred milligram aliquots of feces were weighed, incubated with 2 ml of chloroform-methanol (2:1,v/v) for 30 min at 60°C with constant agitation, and then centrifuged (5 min at 5,000 rpm). Water (1 mL) was added to the supernatant, and following vortex, phase separation was induced by low-speed centrifugation (2,000 rpm for 10 min). The lower chloroform phase was then removed and transferred to a new tube, and the sample was dried under nitrogen. Then samples were re-suspended in 500 μL chloroform, evaporated to dryness, and finally re-suspended in 500μL 100% ethanol. For analysis of liver lipids, 100-mg aliquots of tissue were homogenized with 2 mL chloroform-methanol and then agitated overnight on an orbital shaker at 4°C. The homogenate was then centrifuged (5 min at 5,000 rpm), 0.9% NaCl solution was added to the liquid phase, and the samples were vortexed. Phase separation was induced by centrifugation (2,000 rpm for 10 min), and the bottom phase was removed to a new tube and then processed as above. The levels of TC, TG, and FFA in the fecal and liver lipid extracts were measured using commercial kits following the manufacturer’s instructions and normalized to feces or tissue weights.

### Glucose and insulin tolerance tests

For glucose tolerance tests (GTTs), mice were performed after an overnight fast. Glucose (Sigma, USA) was injected (intraperitoneal [i.p.] injection of a 40% solution, 1.5g/kg body weight), and tail vein blood glucose levels were measured at 0, 30, 60, 90, and 120 minutes after injection with a Onetouch Ultra blood glucose monitoring system (LifeScan, USA). For insulin tolerance tests (ITTs), mice were fasted for 4 h and then intraperitoneally injected with human insulin (0.75 U/kg body weight; NovoRapid, Novo Nordisk). Glucose concentrations in blood were measured after 0, 30, 60, 90, and 120 min. Area under the curves (AUC) for GTT and ITT were calculated by trapezoidal approximation, using GraphPad Prism 6.0.

### RNA isolation and quantitative real-time PCR analysis

Total RNA was extracted by using an RNAiso reagent kit (Takara, Japan). Reverse transcription was performed on each RNA sample (1μg) by using PrimeScript RT reagent kit with gDNA eraser (Takara) with a final reaction volume of 20μL. For qRT-PCR analysis, the reaction mixture was run in an iQ5 real time PCR machine (Bio-Rad Laboratories, Hercules, CA, USA) instrument using SYBR Premix Ex Taq II (Takara, Japan). All expression levels were normalized to the corresponding GAPDH mRNA levels and analyzed using the 2^-ΔΔ^CT^^ method. The qRT-PCR cycles were as follows: Step 1, preparative denaturation (30 s at 95 °C); Step 2, 40 cycles of denaturation (5 s at 95 °C) and annealing (30 s at 60 °C); Step 3, dissociation following the manufacturer’s protocol. PCR primers used in the real-time-PCR analysis are listed in Table S1.

### Semi-quantification PCR

The total RNA was extracted by using an RNAiso reagent kit. The cDNA are synthesized using PrimeScript RT reagent kit. PCR was performed in 20μL reaction mixture according to the instruction of KOD-Plus (TOYOBO, Japan). Primers used for GPR109A were indicated at table 1. PCR was performed using an initial denaturation at 94 °C for 5 minutes, followed by 28 cycles at 94 °C for 30 seconds, 53 °C for 30 seconds, and 68 °C for 30 seconds. After 28 cycles, an additional elongation step was performed at 68 °C for 10 minutes. Amplified PCR products were run on 1.2% agarose gels containing ethidium bromide.

### Histological analysis

To examine hepatic steatosis, liver samples were embedded in Tissue-Tek (SA-KURA, USA), frozen and sectioned (10 μm) and stored at −80 °C until use. The sections were stained with hematoxylin and Oil red O (Sigma Aldrich, USA) for lipid deposition.

For histology, adipose tissues were fixed in 10% neutral-buffered formalin, followed by dehydration with graded ethanol (80-100%) and embedding in paraffin. Then the tissues were sliced into 6μm pieces using microtome, de-paraffinized in xylene, passed through 80-100% ethanol. And hematoxylin and eosin (H&E) staining was performed using the standard techniques.

For transmission electron microscopy (TEM) analysis, liver and muscle tissues were cut into 1 mm3 fragments and fixed by immersion in 2.5% glutaraldehyde in phosphate buffer (0.1 M, pH 7.4) overnight at 4°C. After that, they were washed in 0.1 M phosphate buffer for three times. Then, they were post-fixated with 1% osmium tetroxide (OsO4) in 0.1 M phosphate buffer for 2 hours at room temperature. Subsequently, they were rinsed with 0.1 M phosphate buffer for three times again. Tissues were dehydrated through graded alcohols (30, 50, 70, 80, 90, 95 and 100%) for 30 min. Last they were embedded in Epon 812. Ultrathin sections (70-90 nm in thickness) of Epon-embedded samples were placed on copper grids and stained with uranyl acetate and lead citrate. Sections were examined with a Hitachi model H-7650 electron microscope (Hitachi, Japan).

### Measurement of enzymes activities

To determine the activities of enzymes involving in lipogenesis in liver and white adipose tissue, the enzyme source fraction was prepared as follows. Briefly, a 10% (w/v) homogenate was prepared in phosphate buffered solution (pH 7.2-7.4) and then centrifuged at 2000-3000 rpm for 20 min and the supernatant was transferred into a new tube. Samples were stored at −80 °C until use. All of enzymes, such as pyruvate dehydrogenase (PDH), acetyl CoA carboxylase (ACC), acetyl CoA carboxylase 1 (ACC1), and diacylglycerol acyltransferase 2 (DGAT2), were measure by ELISA (Jin Ma, China).

### Intracellular calcium measurement

The fluorescent Ca^2+^ indicator fura-2 was employed to monitor changes of calcium. HepG2 cells were harvested with a Nonenzymatic Cell Dissociation Solution (M&C Gene Technology, China), washed twice with Hanks’ solution, and resuspended at a density of 5×10^6^ cells/mL in Hanks’ balanced salt solution containing 0.025% bovine serum albumin. The cells were then loaded with 3 μM Fura-2-AM (Dojindo Laboratories) for 30 min at 37 °C. Cells were washed twice in Hanks’ solution, and then resuspended in Hanks’ solution at a concentration of 1×10^7^ cells/mL. Cells were stimulated with the indicated concentrations of niacin and Ca^2+^ flux was measured using excitation wavelengths of 340 and 380 nm in a fluorescence spectrometer.

### Western blot analysis

Proteins were extracted from tissues or cultured cells by homogenizing or scraping in ice-cold RIPA buffer (50mM Tris (pH 7.4), 150mM NaCl, 1% Triton X-100, 1% sodium deoxycholate, 0.1% SDS, 2 mM EDTA) (Beyotime, China) containing 1mM sodium orthovanadate and protease inhibitors (Complete Mini, Roche). Tissue samples were transferred to micro-centrifuge tubes and rotated on a rocking platform for 40 min at 4°C after homogenization on ice. The homogenate was cleared by centrifugation at 4°C for 20 min at 12,000 rpm and the supernatant was transferred into a new tube. Protein concentration in the supernatant was determined using the BCA Protein Assay Kit (Probe Gene, China). Equal amounts of protein samples were subjected to SDS-polyacrylamide gel electrophoresis and transferred to PVDF membranes (Millipore, USA). Membranes were blocked with 5% BSA in TBS containing 0.1% Tween-20 (TBST) for 1 h at room temperature, and incubated overnight at 4°C with primary antibodies against p-ERK1/2 (Cat# 4370), ERK1/2 (Cat# 9102), p-AKT (Ser473) (Cat# 4058), AKT (Cat# 4691), p-AMPKα (Thr 172) (Cat# 2535), AMPKα (Cat# 5832), p-ACC (Ser79) (Cat# 11818), ACC (Cat# 3676), β-tubulin (Cat# 2128) (Cell Signaling Technology, USA), GPR109A (Santa Cruz Biotechnology, USA, Cat# sc-134583) respectively. The following day, the membranes were washed with TBST and incubated with a horseradish peroxidase-conjugated secondary antibody (1: 2,000) for 2 hours at room temperature. Proteins were visualized with an enhanced chemiluminescence (ECL) reagent (Beyotime, China) by the Tanon 5200 Chemiluminescent Imaging System (Tanon, China). The target proteins were quantified by densitometric analysis with Image J software and normalized to β-tubulin level.

### Statistical analysis and data processing

For animal studies, n values indicate the number of animals per cohort. For animal studies, sample sizes were based on previous studies and expertise. All data of cells shown are representative of at least three independent experiments. All results are expressed as mean ± SEM. The GraphPad Prism 6 software (San Diego, CA) was used for statistical analyses. Unpaired two-tailed Student’s t-test was used for comparisons between two groups for all figures. Differences were considered significant when P < 0.05. All P-values for main figures and EV figures can be found in Appendix Tables S2-S12.

## Abrreviations

NA: nicotinic acid;
HFD: high fat diet;
NCD: normal chow diet;
TG: triglyceride;
TC: total cholesterol;
FFA: free fatty acid;
LDL: low-density lipoprotein;
HDL: high density lipoprotein;
WAT: white adipose tissue;
eWAT: epididymal white adipose tissue;
pWAT: perirenal white adipose tissue;
iWAT: inguinal white adipose tissue;
sWAT: subcutaneous white adipose tissue;
BAT: brown adipose tissue;
ITT: insulin tolerance test;
GTT: glucose tolerance test;
PDH: pyruvate dehydrogenase;
ACC: acetyl CoA carboxylase;
ACC1: acetyl CoA carboxylase 1;
DGAT2: diacylglycerol acyltransferase 2;
FAS: fatty acid synthase;
SREBP-1c: sterol regulatory element-binding transcription factor 1;
SCD-1: stearoyl-CoA desaturases-1;
FATP4: fatty acid transport protein 4;
I-FABP: intestinal-type fatty acid-binding protein;
NPC1L1: Niemann-Pick C1-Like 1;
UCP1: uncoupling protein 1;
PGC-1α: peroxisome proliferator-activated receptor gamma coactivator 1-alpha;
PRDM16: PR domain containing 16;
AUC: area under the curves;

